# Activation of the Zinc-sensing receptor GPR39 promotes T cell reconstitution after hematopoietic stem cell transplant

**DOI:** 10.1101/2021.09.02.458741

**Authors:** Lorenzo Iovino, Kirsten Cooper, Paul deRoos, Sinéad Kinsella, Cindy Evandy, Tamas Ugrai, Francesco Mazziotta, Kathleen S Ensbey, David Granadier, Kayla Hopwo, Colton Smith, Alex Gagnon, Sara Galimberti, Mario Petrini, Geoffrey R. Hill, Jarrod A. Dudakov

## Abstract

Prolonged lymphopenia represents a major clinical problem after cytoreductive therapies such as chemotherapy and the conditioning required for hematopoietic stem cell transplant (HCT), contributing toward the risk of infections and malignant relapse. Restoration of T cell immunity is dependent on tissue regeneration in the thymus, the primary site of T cell development; although the capacity of the thymus to repair itself diminishes over lifespan. However, although boosting thymic function and T cell reconstitution is of considerable clinical importance, there are currently no approved therapies for treating lymphopenia. Here we found that Zinc (Zn), is critically important for both normal T cell development as well as repair after acute damage. Accumulated Zn in thymocytes during development was released into the extracellular milieu after HCT conditioning, where it triggered regeneration by stimulating endothelial cell-production of BMP4 via the cell surface receptor GPR39. Dietary supplementation of Zn was sufficient to promote thymic function in a mouse model of allogeneic HCT, including enhancing the number of recent thymic emigrants in circulation; although direct targeting of GPR39 with a small molecule agonist enhanced thymic function without the need for prior Zn accumulation in thymocytes. Together, these findings not only define an important pathway underlying tissue regeneration, but also offer an innovative preclinical approach to treat lymphopenia in HCT recipients.

**KEY POINTS:**

- Thymocytes release zinc after HCT conditioning is sensed by GPR39 and promotes epithelial repair
- Pharmacologic stimulation of GPR39 promotes T cell reconstitution after HCT

## INTRODUCTION

The thymus is a specialized organ responsible for the generation and maintenance of T cells, a major component of the adaptive immune system. T cell development is a complicated process that requires the close interaction between hematopoietic precursors and the thymic stromal microenvironment, which is comprised of thymic epithelial cells (TECs), fibroblasts, and endothelial cells (ECs). These interactions drive the commitment, proliferation, and differentiation of hematopoietic precursors imported from the circulation in a tightly regulated process^1^.

Despite its importance for the generation and maintenance of a diverse naïve T cell repertoire, the thymus is extremely sensitive to injuries such as that caused by infection, shock, or common cancer therapies including cytoreductive chemo- or radiation therapy^2,3^. Although the thymus harbors considerable regenerative capacity, there is continual repair in response to involution due to aging as well as everyday insults such as stress and infection. Profound thymic damage caused by common cancer therapies. Additionally, the conditioning regimens used as part of hematopoietic cell transplantation (HCT) leads to prolonged T cell deficiency; precipitating high morbidity and mortality from opportunistic infections and likely facilitating malignant relapse^4-10^. At the present time there are no clinically approved strategies to enhance post-transplant T cell reconstitution in recipients of HCT.

Zinc (Zn) is the second most abundant trace element in the body that is capable of interacting with more than 300 proteins involved in almost all aspects of cell function^11-13^, including a well-established role in immune health^14-16^. Much of what we know about the effect of Zn on immune function comes from studies where dietary Zn has been deficient, either due to reduced intake as a result of malnourishment, or via genetic means such as loss of function of ZIP4, a Zn transporter, which clinically leads to the condition *Acrodermatitis enteropathica*^17-20^. In all of these settings of Zn deficiency, widespread immune effects can be seen, including defective B cell development, atrophy of the thymus, and disrupted T cell function^14,21-25^. However, while Zn deficiency (ZD) is known to lead to thymic involution, and supplementation with dietary Zn can ameliorate this phenotype^16,26^, the mechanisms by which Zn acts on thymic function is poorly understood.

Here, we have shown that Zn is critically important for T cell development; with mice fed a Zn-deficient diet exhibiting a profound block in the differentiation and expansion of thymocytes, which was reversed with dietary supplementation of Zn in drinking water. Although Zn directly influences T cell development at the level of thymocytes, we found an increase in the level of extracellular Zn after thymic damage, and this translocation of Zn could directly stimulate the production of BMP4 by ECs, which has recently been found to be a critical for endogenous thymic regeneration after acute injury^27^. This putative role for Zn as a damage-associated molecular pattern (DAMP), was mediated by signaling through the cell surface Zn receptor, GPR39. Notably, not only does dietary zinc supplementation enhance T cell reconstitution after allogeneic HCT, but direct pharmacologic stimulation of GPR39 enhances thymic regeneration and abrogate the need for prolonged Zn administration. These studies demonstrate that Zn is not only important for intrathymic T cell maturation, but also that Zn availability finely tunes mechanisms of thymic regeneration. The studies outlined here therefore not only have the potential to define important pathways underlying tissue regeneration but could also result in innovative clinical approaches to enhance T cell reconstitution in recipients of HCT.

## RESULTS

### Zinc is crucial for steady-state T cell development and promoting regeneration after acute damage

To model the effect of Zn on steady state thymic function, mice were fed a Zn deficient (ZD) diet and thymic function was assessed. Mice fed a ZD diet exhibited reduced thymic cellularity (**Fig. 1A, S1A**) in as little as three weeks of ZD treatment when compared to age-matched mice that received normal chow; thymic cellularity and size declined even further by 8 weeks. Notably, these effects were observed even when there were no gross phenotypes such as weight loss (**Fig. S1B**). Although Zn has previously been shown to affect peripheral T cells^14,22,28,29^, there was no effect of the ZD treatment on absolute lymphocyte count at 21 days (**Fig. S1C**), with significant decrease of naïve T cells seen from 5 weeks of ZD diet (**Fig. S1D**). Importantly, levels of cortisol, a stress hormone with considerable negative effects on the thymus ^3^, was consistent throughout the experiment (**Fig. S1E**). Cell depletion was not uniform amongst developing T cells as at day 21 there was a significant decrease only in double positive (DP) and single positive CD4+ and CD8+ (SP4 and SP8) cells, but no change in the earlier double negative (DN), early thymic progenitors (ETP), or intermediate Single Positive (iSP) thymocytes (**Fig. 1B-C, S2**). Within DP thymocytes, there appeared to be a block after ZD treatment that was marked by different level of expression of Thy1 (**Fig. 1D**), and a corresponding decline in proliferation (**Fig. 1E**). Notably, by 56 days after ZD, all thymocyte subsets were depleted with significantly reduced proliferation in all subsets (**Fig. 1E**).

**Figure 1.**
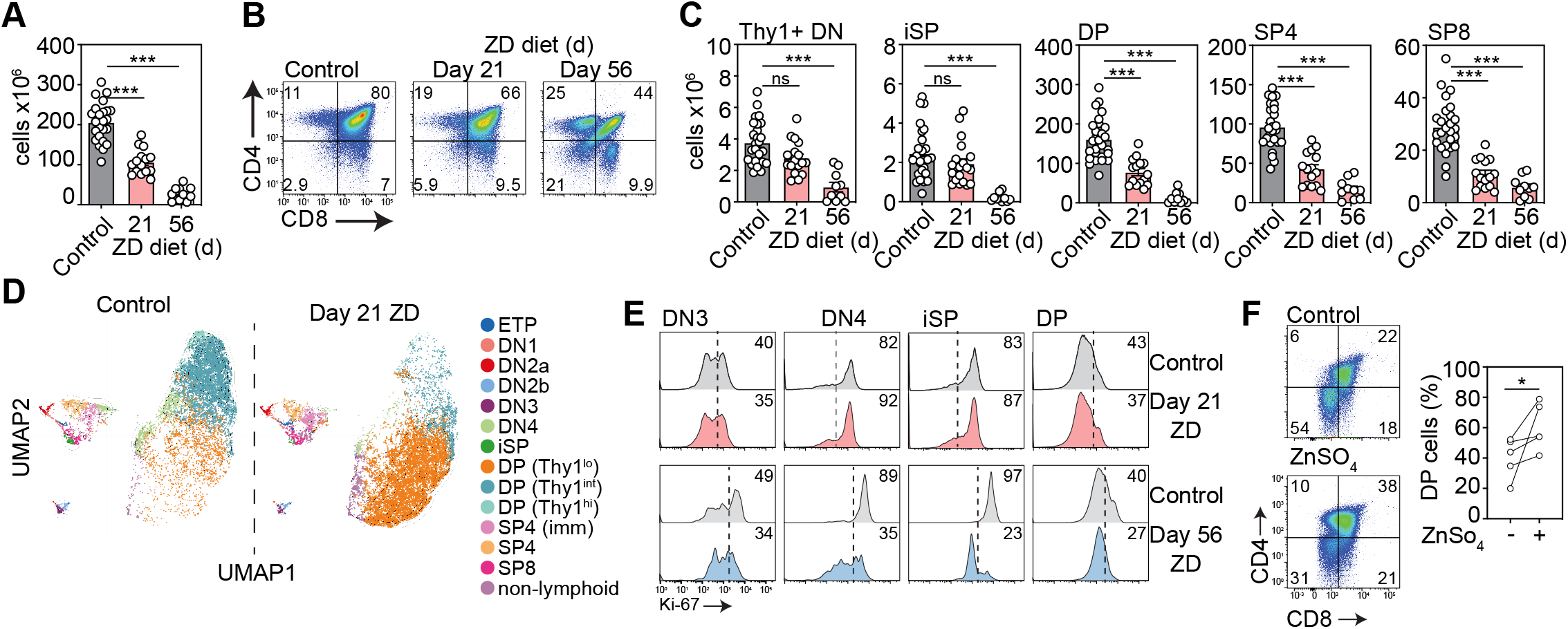
Dietary deficiency of zinc rapidly impairs T cell development. 6-8-week-old female C57BL/6 mice were fed a normal or Zn-deficient (ZD) diet for up to 8 weeks. **A**, Total thymus cellularity from untreated mice or after 21 or 56 days of ZD (untreated, n=24, combined from animals harvested alongside either day 21 or day 56 mice; day 21, n=15 over three independent experiments; day 56, n=10 over two independent experiments). **B**, Concatenated flow cytometry plots displaying CD4 and CD8 expression in the thymus (plots were gated on viable CD45+ cells). **C**, Total number of CD4-CD8-Thy1+ DN, CD3-CD4-CD8+ iSP, CD4+CD8+ DP, CD3+CD8-CD4+ SP4, or CD3+CD8+ CD4-SP8 thymocytes from untreated mice or after 21 or 56 days of ZD (untreated, n=24, combined from animals harvested alongside either day 21 or day 56 mice; day 21, n=15 over three independent experiments; day 56, n=10 over two independent experiments). **D**, Multi parameter flow cytometry data from untreated or 21 days after ZD was placed in UMAP space and clusters were generated based on relative MFI of markers of thymocyte maturation (CD25, CD44, Thy1, CD4, CD8, CD3). **E**, Concatenated flow cytometry plots showing Ki-67 expression in DN3, DN4, iSP, and DP thymocytes from untreated mice or after 21 or 56 days of ZD. **F**, Lineage-depleted bone marrow cells were isolated from untreated 6 week old C57BL/6 mice and co-cultured with OP9-DL1 cells for 30 days in the presence or absence of ZnSO4 (10 µM added from day 0). Concatenated flow cytometry plots displaying CD4 and CD8 expression at day 30 (n=5 independent experiments). Graphs represent mean ± SEM, each dot represents a biologically independent observation. *, p<0.05; **, p<0.01; *** p<0.001.

To confirm the importance of Zn on T cell maturation, we co-cultured non-committed hematopoietic stem cells (HSC) isolated from WT C57CL/6 mice into an OP9/Dll1 system, which is able to promote *in vitro* T cell differentiation of progenitors through the expression of the Notch ligand Dll1 on the surface of a bone-marrow-derived stromal cell line^30,31^; compared to the controls, HSC cultured in media with Zn sulfate at 10 µM showed more robust production of DP (**Fig. 1F**).

Perhaps unsurprisingly given its importance for maintaining thymopoiesis, mice that had been on a ZD diet exhibited significantly worse regeneration following acute damage caused by a sublethal dose of total body irradiation (SL-TBI) (**Fig. 2A)**, as reflected amongst all developing thymocyte subsets (**Fig. S3**) as well as the supporting TEC subsets (**Fig. 2B**). Surprisingly, we found that ZD could have an impact on thymic function even in mice with significant GVHD (**Fig. 2C**). The thymus is extremely sensitive to graft versus host disease (GVHD), even in situations where GVHD may not be detected in classic target organs such as skin, gut or liver^32-34^. Importantly, these thymic effects of a ZD diet could be ameliorated by supplementation of drinking water with Zn sulfate (300mg/Kg/day) beginning on the day of irradiation (**Fig. 2D**). These findings clearly demonstrated that even a short-term reduction in Zn intake has a detrimental impact on thymopoiesis and also has a negative impact on post-damage thymic reconstitution. The early thymocyte damage caused by the ZD treatment seems to affect the late maturation stages between DN and DP; this can also explain why ZD affects also thymic reconstitution after SL-TBI and Zn administration can rescue thymic capacity of regeneration.

**Figure 2.**
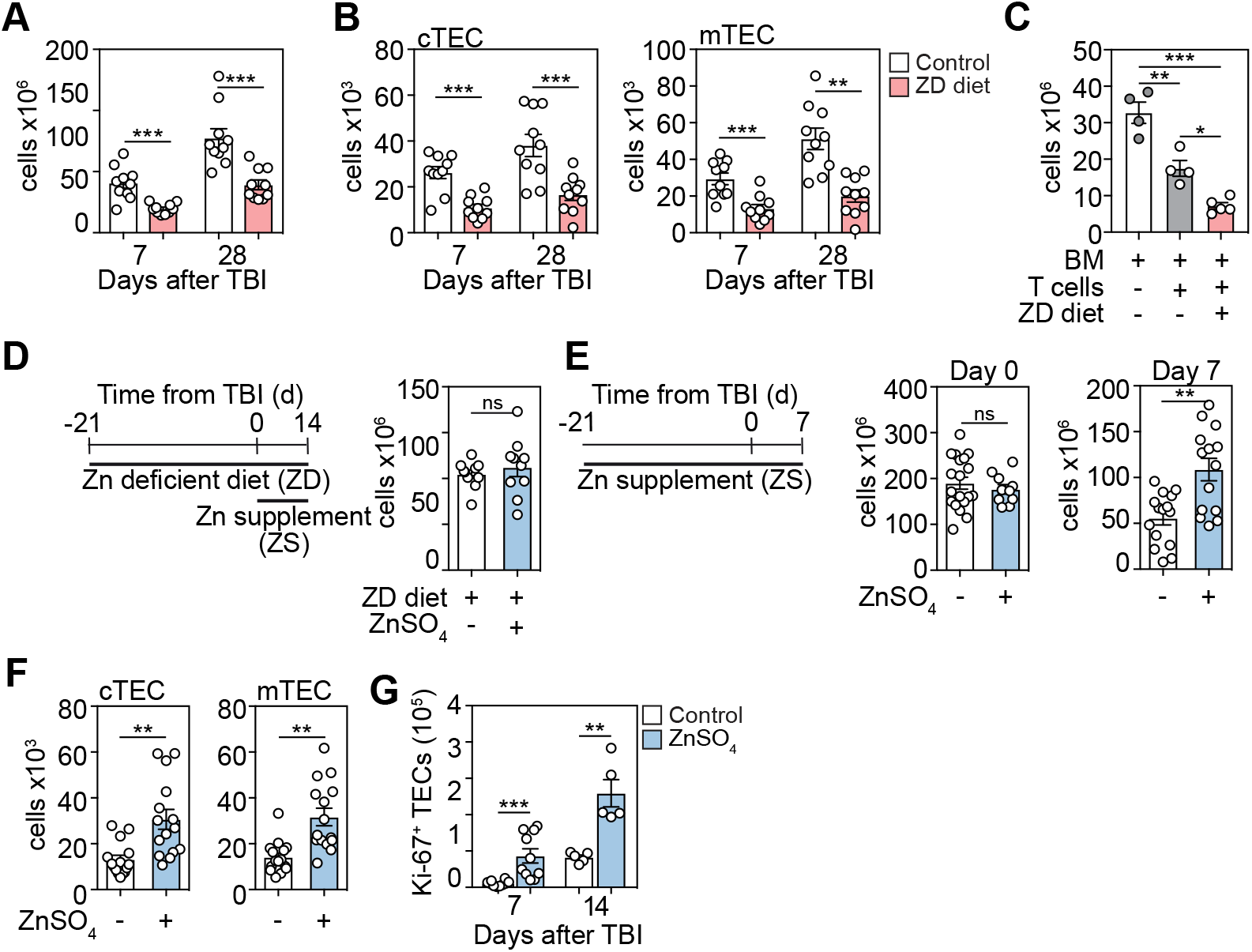
Dietary zinc intake dictates regenerative capacity of the thymus after damage. **A-B**, 6-8 week-old female C57BL/6 mice were fed a normal or ZD diet for 21 days at which point mice were given a sublethal dose of total body irradiation (TBI, 550cGy). **A**, Total thymic cellularity at days 7 and 14 after TBI (n=10/treatment/timepoint across two independent experiments). **B**, Total number of CD45^-^EpCAM^+^MHCII^+^UEA1^lo^Ly51^hi^ (cTECs) and CD45^-^EpCAM^+^MHCII^+^UEA1^hi^Ly51^lo^ (mTECs) at days 7 and 14 after TBI (n=10/treatment/timepoint across two independent experiments). **C**, 6-12 week-old female BALB.B mice were fed with normal or ZD diet for 21 days, at which point mice were given a lethal dose of TBI (900cGy) and 10 × 106 T cell-depleted BM cells from 6-8 week-old C57BL/6. One cohort also received 2 × 10^6^ T cells to induce GVHD; thymic cellularity was quantified on day 14 after allo-HCT (n=5-6/group). **D**, 6-8 week-old female C57BL/6 mice were fed with normal or ZD diet for 21 days at which point mice were given 550 cGy TBI and total thymus cellularity quantified on day 14. One cohort was given supplemental Zn in drinking water (300mg/kg/day ZnSO_4_) from day 0 until analysis on day 14 (n=10/group across two independent experiments). **E-F**, 6-8 week-old female C57BL/6 female mice were given supplemental Zn in drinking water (300mg/kg/day ZnSO_4_) for 21 days at which point mice were given 550 cGy TBI. Mice were maintained on ZnSO4 in drinking water and the thymus was analyzed on day 0 or 7 (Day 0: untreated, n=22; ZD, n=10 across two independent experiments; Day 7: untreated, n=15; ZD, n=15, across three independent experiments). **E**, Total thymic cellularity. **F**, Total number of cTECs and mTECs. **G**, Total number of Ki-67+ TECs. Graphs represent mean ± SEM, each dot represents a biologically independent observation. *, p<0.05; **, p<0.01; *** p<0.001.

Since ZD treatment impairs thymic function and immune recovery after SL-TBI, we hypothesized that dietary Zn supplementation could improve thymic reconstitution after acute insult. Mice were put on Zn supplementation (ZS) (300 mg/Kg/day/mouse of Zn sulfate monohydrate, in drinking water) ^26^ for three weeks prior to SL-TBI and maintained on ZS until day 7 after TBI. Although we observed no difference in thymic cellularity between control and ZS mice after three weeks of treatment, mice that received ZS treatment exhibited improved reconstitution after SL-TBI (**Fig. 2E**), reflected by individual thymocyte populations (**Fig. S4**) and within TEC subsets (**Fig. 2F**). Both the absolute number and the proportion of proliferating TECs, measured by the expression of Ki-67, were higher in the thymuses from mice that received Zn supplementation (**Fig. 2G**).

### Zinc stimulates the production of BMP4 by thymic endothelial cells

Given the increased proliferation of TECs after Zn supplementation, we tested the direct effect of ZnSO_4_ on proliferation of mouse cortical (C9) and medullary (TE-71) TEC cell lines. Using this approach, we did not observe any direct effect of Zn on TECs proliferation, or their ability to express key thymopoietic transcription factors such as FOXN1 (**Fig. S5A-B**). One mechanism by which TECs are induced to proliferate is via stimulation by BMP4, which is produced by ECs in response to damage and can mediate thymic repair by stimulating TEC regeneration^27,35^. Interestingly, there is also a significant body of work demonstrating that Zn plays an important role in vascular integrity and response to stress of ECs^36-38^. To determine if BMP4 could be a mediator of the effect of Zn in thymic regeneration, we first assessed the level of BMP4 in the thymus of mice that had received either a ZD diet for three weeks before TBI (and throughout the study) or mice that had received a ZD diet but had also been given ZS in drinking water. We assessed the expression of BMP4 by ELISA at day 10, the timepoint that we have found previously to be at the peak of expression after damage^27^. We found that indeed mice that had received a ZD diet had significantly reduced levels of BMP4 in the thymus, but dietary supplementation of Zn restored BMP4 levels (**Fig. 3A**). Consistent with these findings, mice that had been given ZS without prior ZD had increased levels of BMP4 (**Fig. 3B**), and purified ECs from ZS-treated mice exhibited increased expression of *Bmp4* measured by qPCR (**Fig. 3C**). Together, these findings suggest that Zn is not only involved in thymocyte maturation, but also in overall thymic regeneration by stimulating the production of BMP4 from EC.

**Figure 3.**
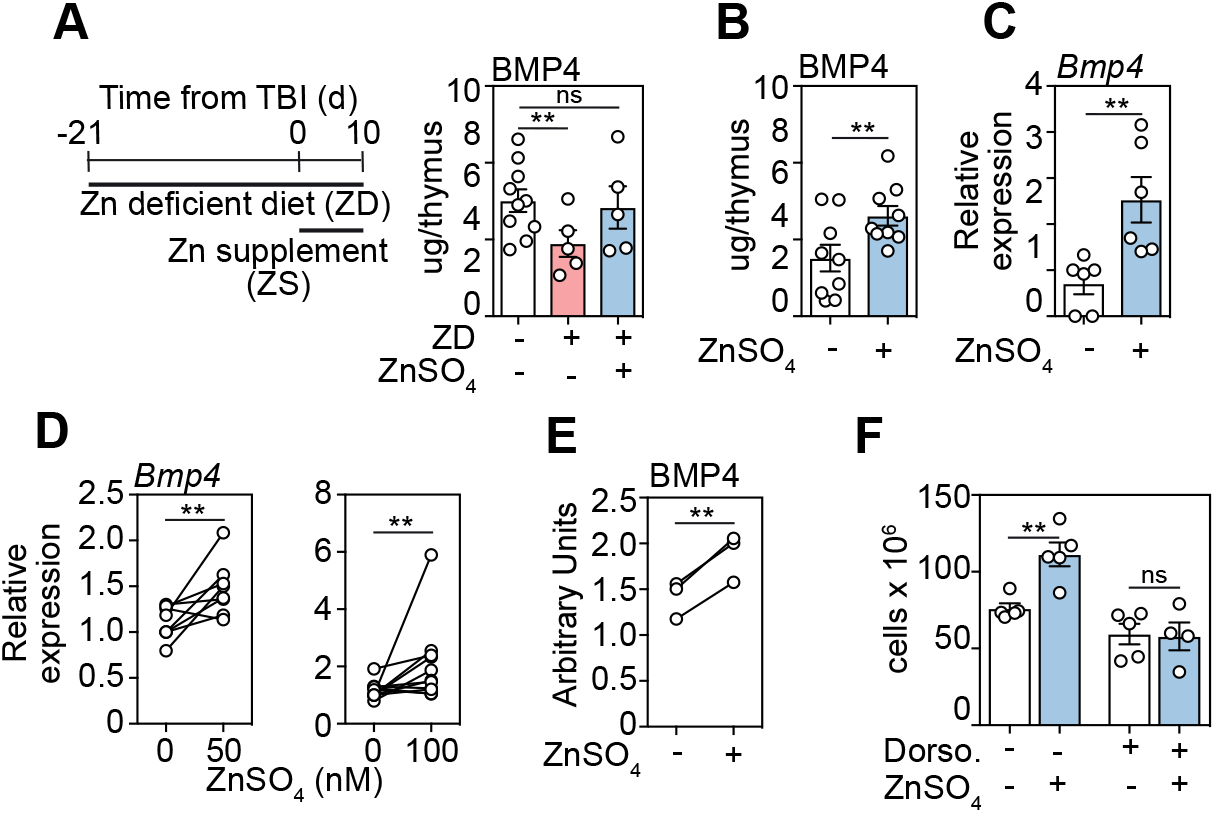
Zinc stimulates the production BMP4 by endothelial cells. **A**, 6-8 week-old female C57BL/6 mice were fed a normal or ZD diet for 21 days at which point mice were given 550 cGy TBI. One cohort was given supplemental Zn in drinking water (300mg/kg/day ZnSO_4_) from day 0. Levels of BMP4 were quantified by ELISA at day 10 after TBI (n=5-10/group from one independent experiment). B-C, 6-8-week-old C57BL/6 female mice were given supplemental Zn in drinking water (300mg/kg/day ZnSO_4_) for 21 days at which point mice were given 550 cGy TBI. Mice were maintained on Zn-supplemented drinking water for the duration of the study. **B**, BMP4 levels measured by ELISA at day 10 (n=9/group combined from three independent experiments). **C**, ECs were FACS purified at day 7 and Bmp4 expression was measured by qPCR (n=6/group combined from two independent experiments). **D-E**, exECs were generated as previously described (27, 39) and stimulated for 24 hours with ZnSO4 at the indicated concentrations and Bmp4 expression measured by qPCR at 24 hours (50µM: n=8/group combined from three independent experiments; 100µM: n=11/group across five independent experiments) **(D)** and BMP4 protein was quantified by ELISA at 48 hours (n=3 independent experiments) **(E). F**, 6-8 week-old female C57BL/6 mice were given supplemental Zn in drinking water (300mg/kg/day ZnSO4) for 21 days at which point mice were given 550cGy TBI. Mice were administered with the BMP type I receptor inhibitor Dorsomorphin dihydrochloride (12.5mg/kg) i.p. at day −1 before TBI and twice daily after TBI and all mice were maintained on ZnSO4 in drinking water for the duration of the study. Total thymus cellularity was quantified at day 10 after TBI (n=5/group from one independent experiment). Graphs represent mean ± SEM, each dot represents a biologically independent observation. *, p<0.05; **, p<0.01; *** p<0.001.

To determine if there was a direct effect of Zn on thymic ECs, we constitutively activated the Akt pathway using the prosurvival adenoviral gene E4ORF1, which allows ECs from multiple tissues, including the thymus, to be propagated and manipulated ex vivo while maintaining their phenotype and vascular tube formation for functional manipulation and *in vitro* modeling of regenerative pathways^27,39,40^. Using this approach, we found that ECs showed a dose-dependent increase in the transcription of *Bmp4* by qPCR after 24 hours of exposure to exogenous Zn (**Fig. 3D**). This finding was confirmed at the protein level after 48 hours of exposure to Zn (**Fig. 3E**). Consistent with our hypothesis that Zn is involved in the *in vivo* pathway of BMP4 production, treatment with the pan-BMP-receptor inhibitor dorsomorphin dihydrochloride abrogated the effect of ZS on thymic regeneration (**Fig. 3F**).

### Zinc translocation into the extracellular milieu after acute damage stimulates endogenous production of BMP4 by ECs

In order to clarify how Zn can mechanistically contribute to endogenous thymic regeneration, we first measured changes in Zn levels in otherwise untreated WT mice after TBI by inductively coupled plasma mass spectrometry (ICP-MS). This analysis revealed that the total amount of Zn in whole thymic lysates (both intracellular and extracellular compartments) decreased after damage, following the same trend as thymic cellularity (**Fig. 4A**)^27^. When we assessed extracellular Zn as a function of total Zn, we found a significant translocation of Zn from intracellular to extracellular space after damage (**Fig. 4B**).

**Figure 4.**
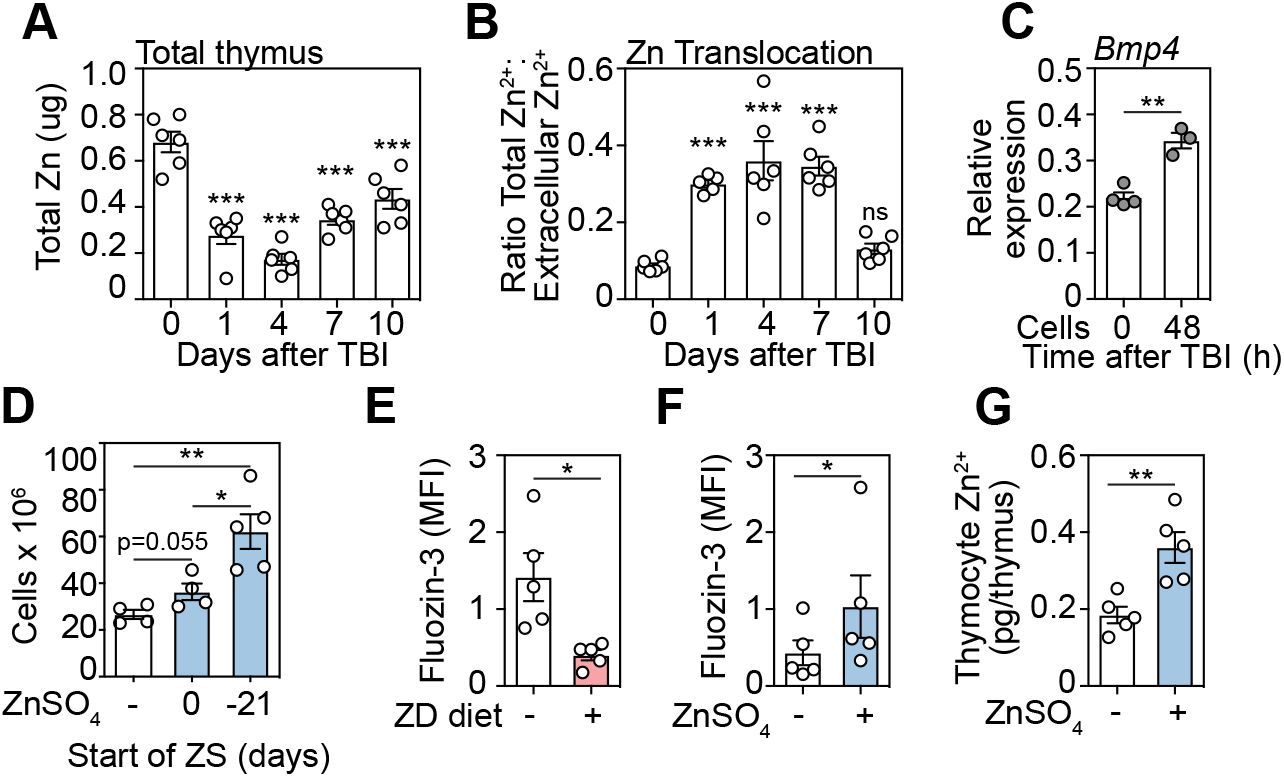
Zn accumulates in thymocytes and is released after damage. **A-B**, 6-8 week-old female C57BL/6 mice were given 550 cGy TBI and levels of Zn were measured by inductively coupled plasma mass spectrometry (ICP-MS). **A**, Total thymic amounts of Zn from both intracellular and extracellular fractions of thymus (n=6/timepoint). **B**, Extracellular Zn was measured only in thymic supernatants and the ratio of extracellular to total thymic Zn was calculated (n=6/timepoint). **C**, 6-8 week-old female C57BL/6 mice were given supplemental Zn in drinking water (300mg/kg/day ZnSO_4_) for 21 days at which point one cohort was given 550cGy TBI. Thymocytes were isolated either before or 48 hours after TBI and co-cultured with exECs. Bmp4 expression was measured by qPCR at 24 hours (n=3-4/group). **D**, 6-8 week-old C57BL/6 female mice were given supplemental Zn in drinking water (300mg/kg/day ZnSO_4_) for either 21 days before TBI, or from the day of TBI and maintained on ZnSO_4_ in drinking water for the duration of the study. Thymus cellularity was measured at day 28 after TBI (n=4-5/group). **E**, 6-8 week-old female C57BL/6 mice were fed a normal or ZD diet for 21 days after which thymocytes were isolated by CD90+ magnetic separation. Intracellular Zn levels were measured by staining with Fluozin-3 and assessed by flow cytometry (n=5/group across two independent experiments). **F-G**, 6-8 week-old female C57BL/6 mice were given supplemental Zn in drinking water (300mg/kg/day ZnSO_4_) for 21 days after which thymocytes were isolated by CD90+ magnetic separation and Zn measured by staining with Fluozin-3 **(F)** or ICP-MS **(G)**.Graphs represent mean ± SEM, each dot represents a biologically independent observation. *, p<0.05; **, p<0.01; *** p<0.001.

To functionally assess this finding, we co-cultured exECs with supernatants isolated from ZS-treated mice at day 0 and 48h after TBI. The supernatant of thymuses harvested 2 days after TBI and incubated with exECs for 24 hours increased the expression of *Bmp4* from ECs when compared to the same experiment performed with supernatant from day 0 (**Fig. 4C**). 98% of thymus cellularity is comprised of thymocytes, which are also the most sensitive cells to damage and the cells that require Zn for their generation and maintenance. Given the fact we observed a better effect if ZS is begun several weeks before TBI (**Fig. 4D**), we hypothesized that thymocytes, which normally internalize Zn during their maturation, accumulate Zn during ZS, which allows for increased bioavailability of extracellular Zn after damage, allowing for the triggering of the regenerative response in ECs. Consistent with this hypothesis, thymocytes isolated from mice given a ZD diet exhibited significantly lower levels of Zn (**Fig. 4E**) and mice given ZS in their drinking water showed significantly increased levels of intracellular Zn (**Fig. 4F-G**).

### Zn signals though the cell surface GPCR GPR39 on endothelial cells to stimulate production of BMP4

There are two main modalities by which Zn can mediate its effect on cells, influx (and efflux) using the ZIP (and ZnT) ion channels, whereby Zn can interact directly with over 300 proteins^41,42^; or via cell surface Zn receptors such as the G-protein coupled receptor GPR39^43,44^. In order to identify the putative modality of Zn, we selectively increased the intracellular Zn concentration of exECs by treating them with the Zn ionophore sodium pyrithione; however, we did not observe any increase of *Bmp4* expression using this approach (**Fig. 5A**), suggesting that binding to a surface receptor is more likely than through Zn internalization. Little expression of GPR39 could be detected on thymocyte populations of otherwise untreated WT mice; however, we found significant expression on non-hematopoietic stromal cells such as TECs, fibroblasts, and ECs (**Fig. 5B, S6A**). Notably, while there was no change in expression within TECs or fibroblasts after damage, we found an increase in expression of GPR39 on ECs (**Fig. 5C, S6B**), suggesting their potential to respond to extracellular Zn after damage is increased. Interestingly, recent reports have demonstrated that many functions of ECs can be regulated by GPR39 signaling^43,44^. GPR39 acts by translating extracellular Zn signals into release of intracellular second messengers such as ERK and calcium release^37,45^. Consistent with this, stimulation of exECs with Zn led to phosphorylation of ERK1/2 (**Fig. 5D**), and when we blocked ERK with the inhibitor FR180204 prior to Zn stimulation, BMP4 production was abrogated (**Fig 5D**). Importantly, demonstrating the functional importance of GPR39 for EC-mediated regeneration, Zn-mediated production of *Bmp4* was abrogated in exECs after silencing of *Gpr39* expression (**Figs. 5E, S6C**). To confirm the functional role of GPR39 in the response of EC to Zn, we treated exECs with the selective GPR39 agonist TC-G1008. Stimulation of exEC with this molecule for 24 hours produced an increase in *Bmp4* expression that was higher than with Zn alone (**Fig. 5F**). *Bmp4* did not further increase by combining Zn and TC-G1008.

**Figure 5.**
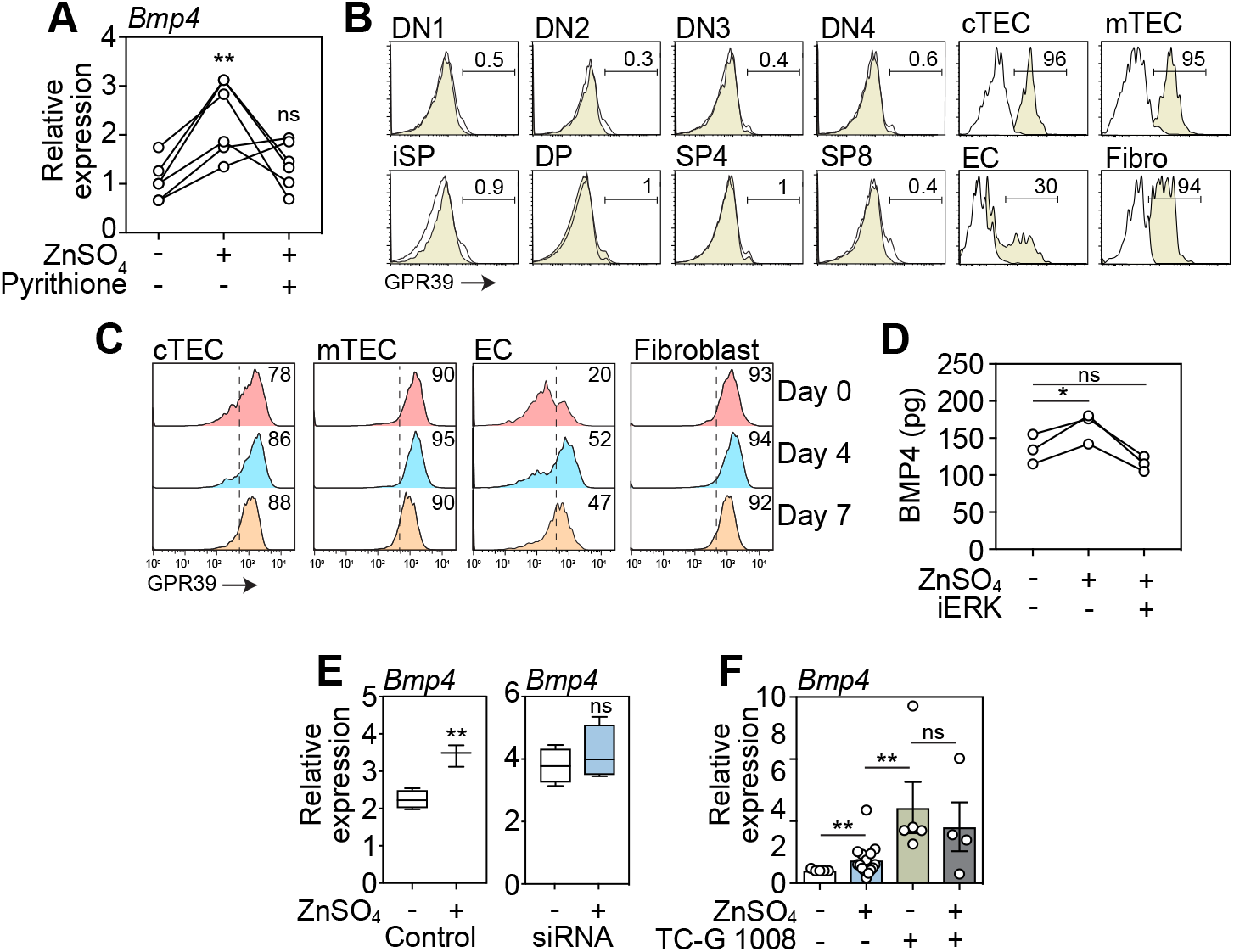
GPR39 expressed by thymic endothelial cells is the central mediator of Zn-centered regeneration. **A**, exECs were stimulated for 24 hours with ZnSO_4_ (100µM) with or without the Zn ionophor pyrythione. Bmp4 expression was measured by qPCR (n=6 combined from two independent experiments). **B**, GPR39 expression across subsets in the thymus by flow cytometry at baseline. Displayed are concatenated plots from one experiment. **C**, Expression of GPR39 on cTECs, mTECs, ECs and Fibroblasts at days 0, 4, and 7 after TBI. Displayed are concatenated plots from one experiment. **D**, exECs were stimulated for 24 hours with ZnSO4 (100µM) with or without the ERK inhibitor FR180204. BMP4 was measured by ELISA. **E**, GPR39 expression was silenced in exECs by siRNA and stimulated for 24 hours with ZnSO_4_ (100µM) after which Bmp4 expression was measured by qPCR (n=3/group). **F**, Bmp4 expression in exEC cultured for 24 hours in presence of ZnSO4 (100μM) and/or the GPR39 agonist TC-G1008 (25μM) (n=5-15 combined from five independent experiments). Graphs represent mean ± SEM, each dot represents a biologically independent observation. *, p<0.05; **, p<0.01; *** p<0.001.

### Zinc supplementation promotes T cell reconstitution after hematopoietic cell transplantation

Thymic regeneration is a particular challenge after the myeloablative conditioning required for successful HSCT^46^. Therefore, we assessed the feasibility of Zn supplementation using a single minor (male)-antigen mismatched model of murine allogeneic T-depleted HSCT where any effects mediated by GVHD are excluded. Similar to our findings in the sublethal TBI model, we found that dietary ZS promoted thymic reconstitution in this setting of severe damage (**Fig. 6A**). Increases were also observed in all developing thymocyte subsets and TEC subsets (**Fig. S7A-B**). In order to track the export of T cells from the thymus, we performed a similar T cell depleted allo-HCT whereby donors expressed green fluorescent protein (GFP) under the control of RAG2, which allows for the detection of cells recently exported from the thymus (referred to as recent thymic emigrants (RTEs)^47,48^. In both peripheral blood and spleen, mice that received ZS showed higher levels of GFP+ CD4+ and CD8+ lymphocytes after HCT (**Fig. 6B**).

**Figure 6.**
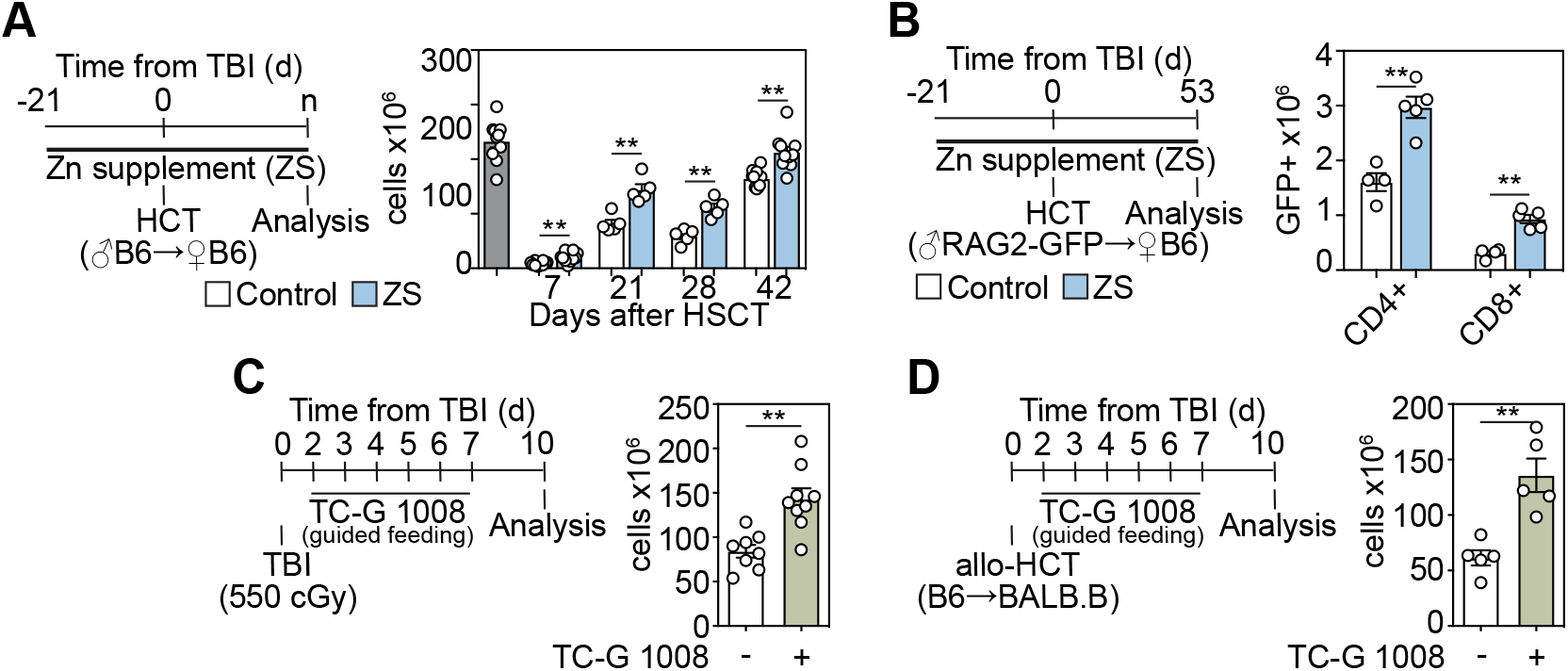
Experimental targeting of the GPR39 receptor improves thymic repair and T cell reconstitution after allo-HCT. **A**, 6-8-week-old male C57BL/6 mice were given supplemental Zn in drinking water (300mg/kg/day ZnSO_4_) for 21 days at which point mice were given a lethal dose of TBI (2 × 550cGy) along with T cell depleted BM from female C57BL/6 mice. Mice were maintained on ZnSO_4_ in drinking water for the duration of the study. Total thymic cellularity at days 7, 21, 28, and 42 after HCT (n=5-10/group/timepoint combined from two independent experiments). **B**, 6-8-week-old male C57BL/6 mice were given supplemental Zn in drinking water (300mg/kg/day ZnSO_4_) for 21 days at which point mice were given a lethal dose of TBI (2 × 550cGy) along with T cell depleted BM from female RAG2-GFP mice. Splenic T cells were analyzed for GFP expression on day 53 (n=4-5/group). **C**, 6-8-week-old C57BL/6 female mice were given 550cGy TBI and either vehicle or TC-G 1008 (20mg/mouse/day) by guided feeding daily from day 0 until day 10 when thymus cellularity was assessed (n=8-9 combined from two independent experiments). **D**, 6-8-week-old female BALB.B mice were given a lethal dose of TBI (900cGy) along with 10 × 106 T cell depleted BM from female C57BL/6 mice and either vehicle or TC-G 1008 (20mg/mouse/day) by guided feeding daily from day 0 until day 8, then on day 10 and 12. Thymuses were harvested and analyzed at day 14 (n=5/group). Graphs represent mean ± SEM, each dot represents a biologically independent observation. *, p<0.05; **, p<0.01; *** p<0.001.

The complicated mechanism by which we believe dietary Zn supplementation promotes thymic regeneration involves a significant period before transplant in order for thymocytes to accumulate Zn to be released after injury, thereby allowing signaling through GPR39 in regeneration-initiating ECs. To test if directly stimulating GPR39 could abrogate this lead time, we treated mice with the GPR39 agonist TC-G1008^49^. Using this approach, we could show that mice treated with TC-G1008 beginning on day 1 after TBI showed significantly improved thymic cellularity (**Fig. 6C**). Moreover, also mice that had been given a T cell depleted allo-HSCT across multiple minor histocompatibility antigens also exhibited increased thymic cellularity following TC-G1008 treatment (**Fig. 6D**). Taken together, these data suggest that improved thymic regeneration caused by Zn signaling can help immune reconstitution after HSCT by increasing the production of thymic-derived naïve T cells. Additionally, this pathway can be pharmacologically targeted by stimulating GPR39 signaling.

## DISCUSSION

Alterations in Zn uptake, retention, sequestration, or secretion can quickly lead to ZD and affect Zn-dependent functions in virtually all tissues, and in particular in the immune system, including thymic involution^17,24,25^. We show that even short-term zinc deprivation in young animals has a profound impact on thymic function, before effects on peripheral immune cells can be detected. ZD reduced the replicative ability of thymocytes, especially in the transition to DP thymocytes. Although the specific mechanism by which Zn is acting on thymocyte development is unclear, there is evidence that many Zn-finger transcriptional factors that heavily depend on general Zn availability are crucial for T cell development^50-52^. In our data, intracellular Zn levels in thymocytes responded rapidly to changes in systemic Zn availability; with lower levels of Zn in thymocytes under ZD and higher levels after ZS. This is consistent with the notion that T cells actively internalize Zn during activation and replication through the expression of Zn importers such as ZIP6^22^. Notably, the changes observed in thymic function preceded the other classic signs of ZD, such as weight loss and skin and fur changes, confirming the sensitivity of thymopoiesis to Zn level; although the contribution of systemic effects cannot be completely ruled out^53^.

Thymic regeneration is a complex and poorly understood process, in which the cytokines produced from damage-resistant cells, such as IL-22 from innate lymphoid cells (ILC), IL-23 from dendritic cells (DC), and BMP4 from endothelial cells (EC), stimulate TECs to proliferate and mediate broader thymic repair^27,54^. BMP4 is a member of bone morphogenic proteins, a family of peptides involved in embryogenesis and homeostasis of many tissues^55^, including thymic organogenesis and maintenance of *Foxn1* expression in TECs^35,56,57^. We found that modulating levels of Zn in the thymus using either ZD or ZS had a concomitant effect on BMP4 expression. Furthermore, stimulation of thymic ECs with ZnSO_4_ directly induced the production of BMP4 in a GPR39-dependent manner; and the administration of a BMP-receptor inhibitor abrogated the effect of ZS on thymic repair. Notably, a similar role for Zn release into the extracellular space after acute damage has been demonstrated to be involved in tissue repair in tissues such as skin and gut^18,58-60^.

The G-protein coupled receptor GPR39 was recently discovered as a “Zn sensing receptor”^37^ with putative roles in tissue repair in the gut and skin^61,62^. However, while Zn is involved in epithelial cell function in other organs^63-65^, and TECs do express GPR39, our data suggest that its role on TECs regeneration after acute injury is likely indirect through BMP4; although we cannot exclude the possibility that GPR39 also mediates effects directly on other stromal cells. Therefore, we can conclude that Zn is not only needed for thymocyte maturation, but also for thymic repair after acute damage by stimulating the production of BMP4 by ECs.

Thymic regeneration is important following myeloablative conditioning required for successful HCT, after which there is prolonged suppression of T cell immunity. The importance of finding strategies to stimulate thymic-dependent immune reconstitution is highlighted by the correlation between RTEs and clinical outcomes following HCT^2,10^. Given its effects on T cell development and on the induction of regenerative factors, it was perhaps not surprising that ZD mice exhibited worse repair following TBI, however, the fact that ZD led to even worse recovery in mice that had fulminant GVHD highlighted its importance for restoration of thymic function after acute damage and highlights the potential for clinically targeting this pathway. Importantly, we could show not only improved thymic function after allogeneic HCT, but also that this enhanced repair was translated into the circulation with increased numbers of RTEs. However, our findings suggest that the therapeutic benefit of dietary Zn supplementation demands an extended pre-treatment in order for thymocytes to accumulate Zn. Therefore, our findings suggesting that directly targeting the GPR39 receptor itself with a pharmacological agonist is an attractive alternative to induce an equivalent reparative response when given at the time of myeloablative conditioning.

In conclusion, these findings highlight the importance of Zn in steady-state T cell development and reveal a role for Zn in endogenous tissue repair. The studies outlined here not only define important pathways underlying tissue regeneration but could also result in innovative clinical approaches to enhance T cell reconstitution in recipients of HCT.

## METHODS

### Mice

4-6 week-old male or female C57BL/6 (CD45.2) or B6.SJL-Ptprca Pepcb/BoyJ (CD45.1) mice were obtained from the Jackson Laboratories (Bar Harbor, USA). RAG2-GFP mice were kindly provided by Dr. Pamela Fink^47^. Custom-made diets (1 ppm Zinc compared to 35 ppm of control diet) were purchased from Labdiet (St. Louis, MO). Zn supplementation was administered orally by dissolving ultrapure Zn sulfate monohydrate (Alfa Aesar, Haverhill, MA) in drinking water (1.06 g/ml, which delivered 300 mg/Kg/mouse/day based on average water consumption).

Sublethal TBI was given at a dose of 550 cGy while lethal TBI was given for HCT experiments (2 × 550 cGy) with no hematopoietic rescue. HCT mice received 5 × 10^6^ T cell depleted BM cells/recipient. Lethal TBI for the mismatched HSCT (C57BL/6 female donors to BALB.b female recipients) was 900cGy total (2 doses of 450cGy) with 10 × 10^6^ T cell depleted BM cells/recipient. In the ZD+GVHD model, 2 × 10^6^ T cells/recipient were infused together with the BM^66^. T cell depletion and positive selection were performed with CD3 biotinylated and streptavidin-coated beads (Miltenyi Biotech, Germany. #130-094-973). All TBI experiments were performed with a Cs-137 γ-radiation source.

For the administration of TC-G 1008 (Tocris, Bristol, UK.), we followed a micropipette-guided drug administration protocol^67^. The vehicle consisted of condensed milk diluted with ultrapure water 1:3. Mice received 20mg TC-G 1008 daily for the indicated duration. The BMP type I receptor inhibitor dorsomorphin dihydrochloride was given to indicated mice at a dose of 12.5mg/kg one day before SL-TBI and then twice daily from day 1. Mice were maintained at Fred Hutchinson Cancer Research Center (Seattle, WA). Animals were allowed to acclimatize for at least 2 days before experimentation, which was performed according to Institutional Animal Care and Use Committee guidelines.

### Cell Isolation

Individual or pooled single cell suspensions of freshly dissected thymuses were obtained and either mechanically suspended or enzymatically digested as previously described. Cell counts were performed on the Z2 particle counter (Beckman Coulter, Pasadena, CA), on Spark 10M chip reader (Tecan, Switzerland) or on a hemocytometer. CD45− cells were enriched by magnetic bead separation using LS columns and CD45 beads (Miltenyi Biotech, Germany. # 130-052-301). Cell suspensions from the spleen were prepared by tissue disruption with glass slides and filtered through a 40 μm filter. Peripheral blood samples were obtained from mice anesthetized with isoflurane, and a drop of 0.5% proparacaine hydrochloride ophthalmic solution (Bausch & Lomb, Tampa, FL) was applied optically 5 minutes prior to sampling. Blood was collected into EDTA capillary pipettes (Drummound Scientific, Broomall, PA). An aliquot of 20μl from each sample was cryopreserved for mass spectrometry analysis (see below). Peripheral blood counts were performed on Element Ht5 automatic counter (Heska, Loveland, CO).

### Cell cultures

Ex-vivo propagated endothelial cells (exEC) were generated as previously described^39^. Cells were cultured in presence of ultrapure Zn sulfate monohydrate purchased from Alfa Aesar (Haverhill, MA. #1113809) at 5μM, 10μM, 25μM, 50μM, 100 μM for 24 hours or 1 nM, 2 nM, 5 nM, and 10 nM in presence of 50 μM sodium pyrythione (Sigma Aldrich, St. Louis, MO. #H3261-1G). TC-G1008 was used in cell cultures at a final concentration of 25μM. Silencing of the zinc receptor Gpr39 was performed by electroporation using Nucleofector electroporation kit (VPI-1001, Lonza) for exECs (Program M-003, Nucleofector 2b, Lonza), and using the GPR39 siRNA Silencer Select purchased from Thermo, Waltham, MA (#4390771). C9 and TE-71: Mouse C9 (cTEC) and TE-71 (mTEC) cells were kindly provided by A. Farr, University of Washington.

### Co-culture experiments

Thymic extracellular fractions, also referred to as “supernatants” (SN) were obtained by mechanically dissociating thymic tissue in defined volumes of PBS buffer. Thymi were collected from control and Zn-supplemented mice, at day 0 and day 2 after SL-TBI. OP9-DL1 cells were kindly provided by J.C. Zuniga-Pflucker, University of Toronto and cultured as previously described^31^ using lineage-negative BM (using a lineage depletion kit, Miltenyi Biotech, Germany). Cells were transferred every 4-5 days on a fresh layer of OP9-DL1 and flow analyzed every week from day 15 of culture to day 35. Flt-3L and IL-7 were purchased from Peprotech (Rocky Hill, NJ).

### ELISA and Western Blot

For detection of BMP4, whole thymus lysates were prepared by homogenizing tissue in RIPA buffer (50 mM Tris pH 7.6, 150 mM NaCl, 1% NP-40, 1% SDS, 0.01% sodium deoxycholate, 0.5 mM EDTA, and protease inhibitors (Thermo, Waltham, MA. #A32955). ExEC were harvested after 48 hours of exposure to Zn and washed with 4C PBS prior to lysing with RIPA buffer. The resulting samples were normalized for total protein by BCA assay (Thermo, Waltham, MA. #23227) and BMP4 levels quantified using the BMP4 ELISA kit (LSBio, Seattle, WA) and read on a Spark 10M plate reader (Tecan, Switzerland). Cortisol levels were measured with ELISA on peripheral blood (R&D systems, Minneapolis, MN). The proteins were resolved on 12% SDS-PAGE gels, transferred onto PVDF membranes (Bio Rad. Hercules, CA). Blots were analyzed using the ECL detection system or scanned with an Odyssey Infrared Imager (LI-COR Biosciences, Lincoln, NE, USA). In vitro cell proliferation of C9, TE-71 and exEC were measured using the CellTiter Non-Radioactive Cell Proliferation Assay (Promega, Madison, WI).

### Flow cytometry, multidimensional analyses, and FACS sorting

For flow cytometry and cell sorting, surface antibodies against CD45 (30-F11), CD31 (390 or MEC13.3), CD90.2 (30-H12), TER-119 (TER-119), CD4 (RM4-5 or GK1.5), CD8 (53-6.7), TCRβ (H57-597), CD3 (145-2C11), CD44 (IM7), CD25 (PC 61.5), CD62L (MEL14), MHC-II IA/IE (M5/114.15.2), EpCAM (G8.8), Ly51 (6C3), CD11c (HL3), IL-7Rα (A7R34), CCR6 (140706), CD45.1 (A20), CD45.2 (104), ki-67 (16A8), and PDGFRα (APA5) were purchased from BD Biosciences (Franklin Lakes, NJ), BioLegend (San Diego, CA) or eBioscience (San Diego, CA). Ulex europaeus agglutinin 1 (UEA-1), conjugated to FITC or Biotin, was purchased from Vector Laboratories (Burlingame, CA). GPR39 conjugated to FITC was polyclonal and purchased from Signalway Antibody (College Park, MD). For spleen and peripheral blood samples, erythrocyte lysis with ACK buffer (Thermo Scientific, Waltham, MA) was performed before staining. Flow cytometric analysis was performed on a Fortessa X50 (BD Biosciences, Franklin Lakes, NJ) and cells were sorted on an Aria II (BD Biosciences) using FACSDiva (BD Biosciences, Franklin Lakes, NJ). For all assays requiring analysis of intracellular cytokines or phosphoproteins, cells were fixed and permeabilized by using Fix Buffer I and Phospho-Perm Buffer III purchased from BD Bioscience (Franklin Lakes, NJ). Fluozin-3 AM was purchased from Thermo Fisher (Waltham, MA). Analysis of flow cytometry experiments was performed on FlowJo (Treestar Software, Ashland, OR). After standard preprocessing to remove dead cells, gated CD45+ T cells were exported in R (version 4.0.2) for further analyses performed with a custom-made script based on Nowicka M et al. workflow (*71*).

### PCR and Microarray

Reverse transcription-PCR was performed with iScript Clear gDNA cDNA synthesis kit (Bio Rad, Hercules, CA. # 1725035). PCR was done on CFX96 (Bio Rad, Hercules, CA) with iTaq Universal SYBR Green (Bio Rad). Relative amounts of mRNA were calculated by the comparative ΔC(t) method or as relative expression. SYBR Green gene expression assays for qPCR, including *Bmp4* (Mm00432087_m1), *Foxn1* (Mm00433948_m1), *beta-actin, Gpr39, Top1* were all purchased from Life Technologies (Carlsbad, CA) and Bio Rad (Hercules, CA). *b-actin*: synthesized by IDT, F: 5’-CACTGTCGAGTCGCGTCC-3’, R: 5’-TCATCCATGGCGAACTGGTG-3’. *Top1*: synthesized by IDT, sequences provided by PrimerBank, PrimerBank ID 6678399a1. *Bmp4*: Biorad qMmuCED0046239. *Foxn1*: Biorad qMmuCED0044924. Microarray data from CD45^-^ cells was analyzed on day 0, 4 and 7 as previously described ^27^ (GSE106982) and differential gene expression of Gpr39 was calculated.

### ICP-MS

Thymuses were harvested, weighed, and then immediately frozen. Supernatants were generated by mechanical dissociation of the thymuses and subsequent separation of the cellular fraction from the non-cellular fraction. Thymocytes were collected by mechanical dissociation of the thymuses and separation of the stromal component. Each sample was added to trace meal clean plastic vials that had been previously acid leached and rinsed several times with high purity 18 MOhm water. Digestion solution (1:1 v/v mix of 50% HNO3 and 10 % H_2_O_2_) was then added and digested in an all plastic and trace metal clean laminar fume hood. All samples were analyzed at a 10x dilution on an ICAP RQ ICP-MS (University of Washington, TraceLab) in 2% optima grade nitric acid. Three isotopes of Zn were analyzed and cross referenced for isobaric interferences. Zn-66 was chosen as the preferred isotope because it exhibited the largest signal with the least amount of interferences. Exact masses of the initial sample, final sample, and each aliquot were taken into account to determine concentrations in units of mg Zn/g thymic tissue or mg Zn/g supernatant. A rhodium internal standard was used to correct for changes in plasma ionization efficiency. Several external standards including USGS T231 were used to ensure accuracy and traceability.

### Statistics

Statistical analysis between two groups was performed with the nonparametric, unpaired Mann-Whitney U test. Statistical comparison between 3 or more groups was performed with the nonparametric, unpaired Kruskall-Wallis test. All statistics were calculated and display graphs were generated in Graphpad Prism. For multiple comparisons, we used One-Way ANOVA with Tukey’s test.

## Acknowledgments

We gratefully acknowledge Eugenio Mocchegiani, Marco Malavolta, and Robertina Giacconi, of INRCA (Istituto Nazionale di Ricovero e Cura per Anziani), Ancona, for their feedback about this project; the Flow Cytometry and Comparative Medicine Shared Resources at the Fred Hutchinson Cancer Research Center; and the TraceLab in the School of Oceanography at the University of Washington, for ICP-MS. We also gratefully acknowledge Juan-Carlos Zuniga-Plucker (University of Toronto) for the gift of OP9-DL1 cells. This study was supported by National Institutes of Health award numbers R01-HL145276 (J.A.D.), Project 2 of P01-AG052359 (J.A.D.), and the NCI Cancer Center Support Grant P30-CA015704. Support was also received from a Scholar Award from the American Society of Hematology (J.A.D.), The Rotary Foundation through a Global Grant (L.I), and the Immunotherapy Integrated Research Center at the Fred Hutch (L.I.).

## Authorship

Contribution: M.P, L.I. and J.D conceived of the idea of this manuscript. JD, and L.I. designed, analyzed and interpreted experiments, and drafted the manuscript; K.C., P.d.R, S.K., C.E., D.G. C.S., K.H. performed experiments; A.G. and T.U. performed ICP-MS; F.M. performed multidimensional analyses of flow cytometry data. G.R.H. and K.S.E. helped design, perform, and interpret the GVHD experiments. S.G. helped revising the manuscript. All authors contributed to the article and approved the submitted version.

---

**Supplementary Figure 1.**
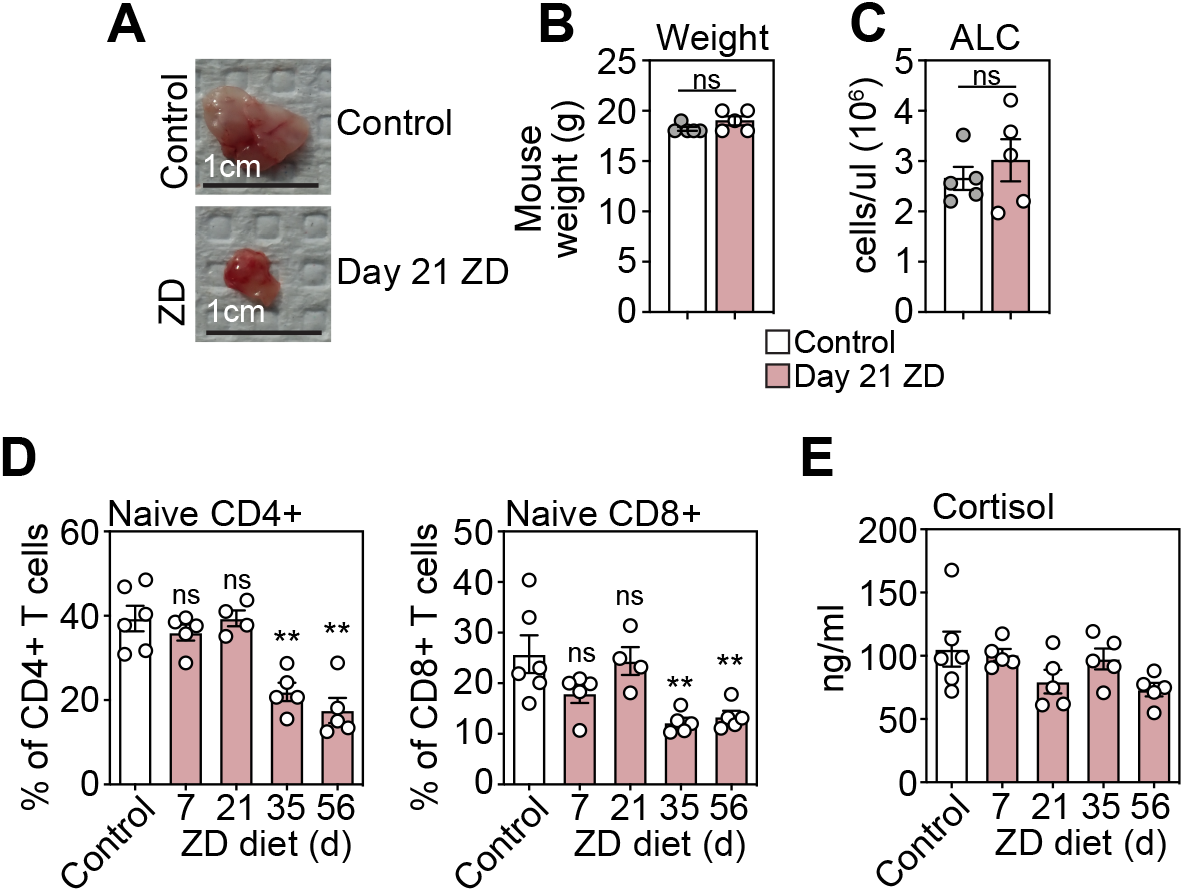
6-8-week-old female C57BL/6 mice were fed a normal or Zn-deficient (ZD) diet for up to 56 days. **A**, Photos of the thymus from mice fed either control diet or ZD diet for 21 days. **B**, Weight of thymuses isolated after 21 days of ZD diet. **C**, Absolute lymphocyte counts (ALC) on the peripheral blood after 21 days of ZD diet. **D**, Proportion of naïve CD4^+^ or CD8^+^ T cells (as a proportion of total CD4+ or CD8+ T cells) after 1, 3, 5, or 8 weeks of ZD diet. **E**, Concentration of cortisol in serum of mice that had received control diet (ctrl), and after 1, 3, 5, and 8 weeks of ZD diet. Graphs represent mean ± SEM, each dot represents a biologically independent observation. *, p<0.05; **, p<0.01; *** p<0.001

**Supplementary Figure 2.**
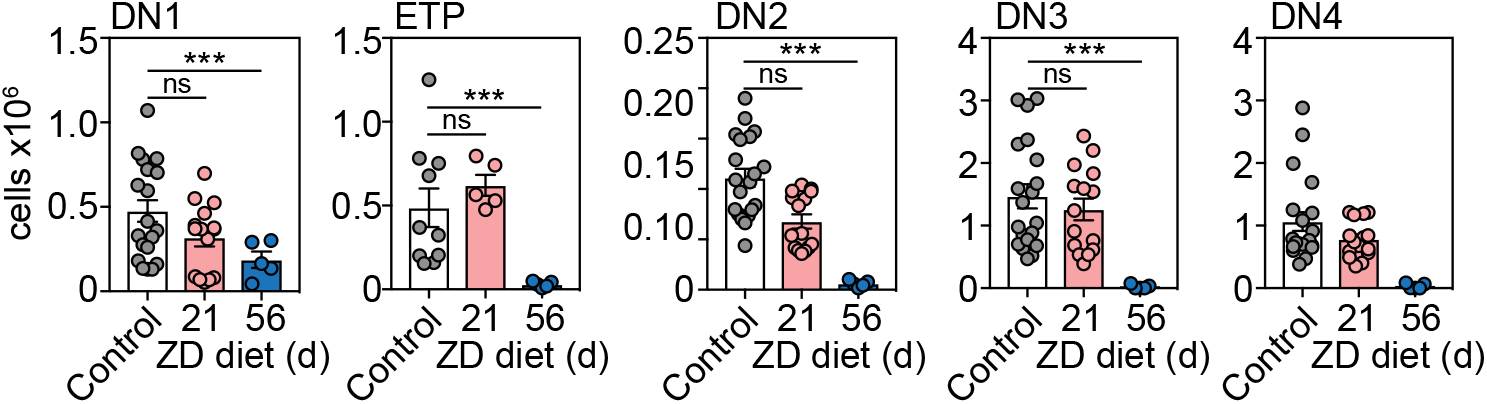
6-8-week-old female C57BL/6 mice were fed a normal or Zn-deficient (ZD) diet for up to 8 weeks. Total number of CD4^-^CD8^-^CD3^-^ double negative thymocytes (DN) from untreated mice or after 21 or 56 days of ZD: CD44^+^CD25^-^ DN1, CD44^+^CD25^-^c-kit^+^ ETP, CD44^+^CD25^+^ DN3, or CD44^-^CD25^-^CD90^+^ DN4 (untreated, n=24, combined from animals harvested alongside either day 21 or day 56 mice; day 21, n=15 over three independent experiments; day 56, n=10 over two independent experiments). Graphs represent mean ± SEM, each dot represents a biologically independent observation. *, p<0.05; **, p<0.01; *** p<0.001.

**Supplementary Figure 3.**
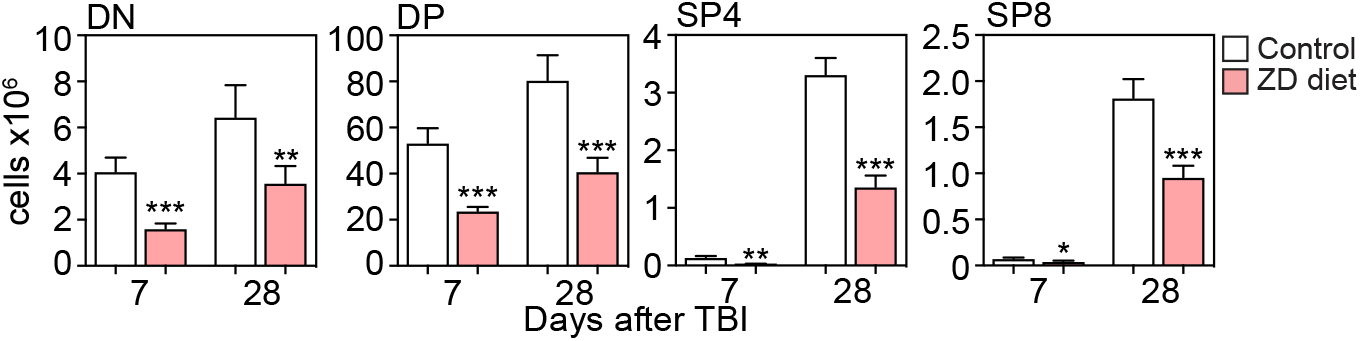
6-8 week-old female C57BL/6 mice were fed a normal or ZD diet for 21 days at which point mice were given a sublethal dose of total body irradiation (TBI, 550cGy). Absolute number of DN, DP, SP4, and SP8 thymocyte subsets was calculated on day 7 and day 28 after TBI. Graphs represent mean ± SEM, each dot represents a biologically independent observation. *, p<0.05; **, p<0.01; *** p<0.001.

**Supplementary Figure 4.**
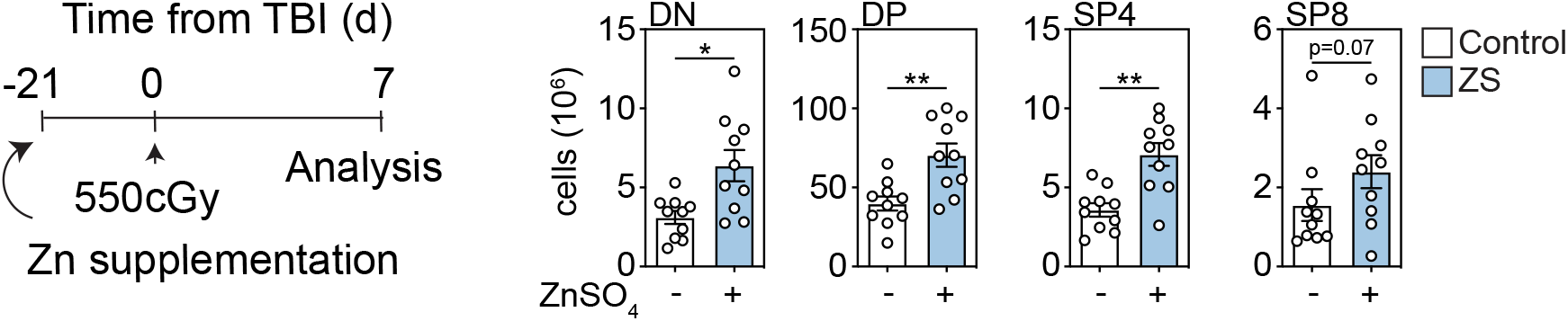
6-8-week-old C57BL/6 female mice were given supplemental Zn in drinking water (300mg/kg/day ZnSO_4_) for 21 days at which point mice were given 550 cGy TBI. Mice were maintained on Zn-supplemented drinking water for the duration of the study. Absolute number of DN, DP, SP4, and SP8 thymocyte subsets was calculated on day 7 after TBI. Graphs represent mean ± SEM, each dot represents a biologically independent observation. *, p<0.05; **, p<0.01; *** p<0.001.

**Supplementary Figure 5.**
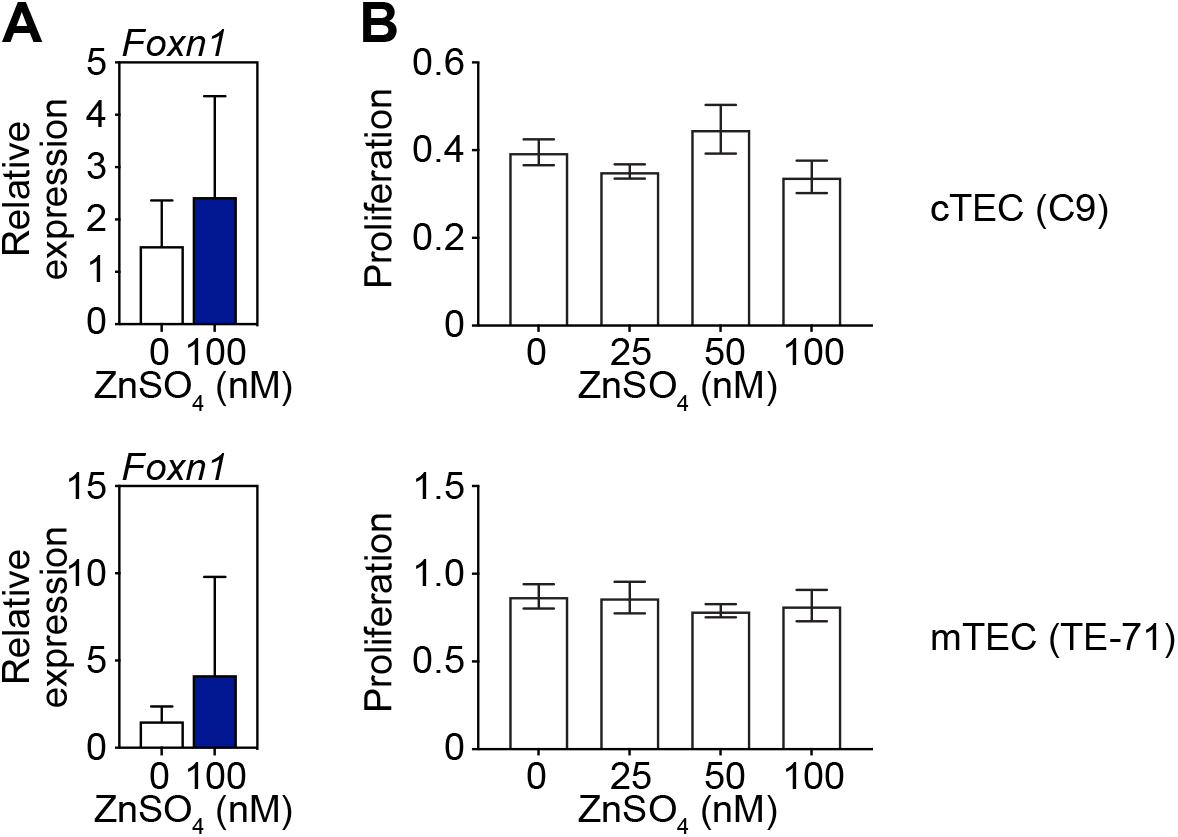
**A**, Thymic epithelial cell lines (C9, cTEC; TE-71, mTEC) were cultured with 100μM ZnSO4 for 24h when Foxn1 expression was quantified by qPCR (n=3 independent experiments). **B**, C9 or TE-71 cells were incubatyed with graded doses of ZnSO4 for 24h after which proliferation was assessed. Graphs represent mean ± SEM, each dot represents a biologically independent observation (n=3 independent experiments). *, p<0.05; **, p<0.01; *** p<0.001.

**Supplementary Figure 6.**
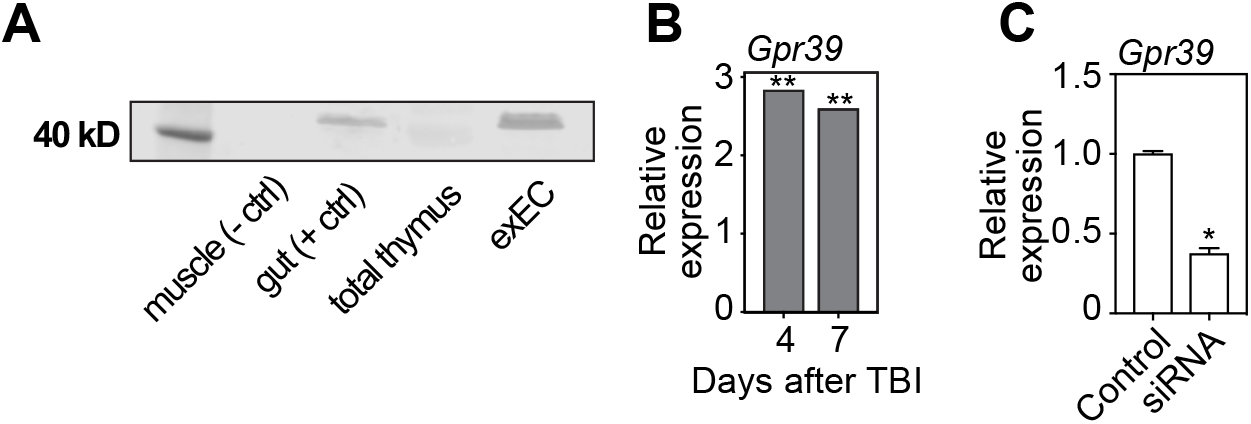
**A**, Western blot showing expression of GPR39 on whole thymus tissue and in thymic exECs. Skeletal muscle was used as negative control and intestine as positive control. **B**, Thymic non-hematopoietic stromal cells were isolated from 6-week-old female C57BL/6 mice using CD45 MACS cell depletion at days 0, 4, and 7 after a single dose sub-lethal total body radiation (SL-TBI) and microarray analysis performed as previously described (Wertheimer et al., 2018) (GSE106982). Displayed is the differential gene expression fold-change of Gpr39 comparing day 4 to day 0 or day 7 to day (n=3; CD45-cells were pooled from 3-4 mice/n). **C**, Gpr39 expression measured by qPCR in exECs after silencing with. Graphs represent mean ± SEM.

**Supplementary Figure 7.**
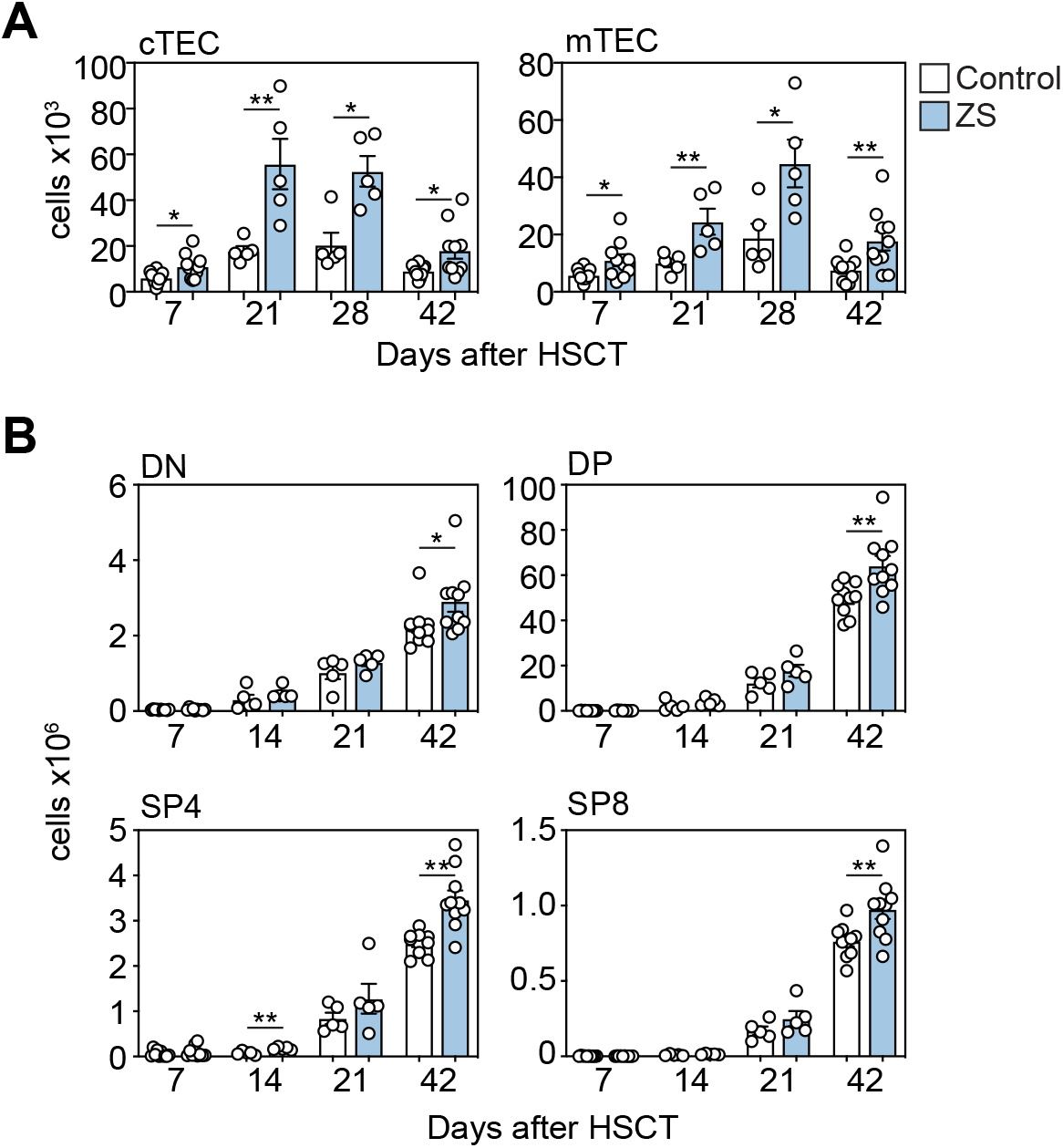
6-8-week-old male C57BL/6 mice were given supplemental Zn in drinking water (300mg/kg/day ZnSO_4_) for 21 days at which point mice were given a lethal dose of TBI (2 × 550cGy) along with T cell depleted BM from female C57BL/6 mice. Mice were maintained on ZnSO4 in drinking water for the duration of the study (n=5-10/group/-timepoint combined from two independent experiments). **A**, Total number of cTECs and mTECs at days 7, 21, 28, and 42 after allo-HCT. **B**, Total number of DN, DP, SP4, and SP8 thymocytes at days 7, 21, 28, and 42 after allo-HCT. Graphs represent mean ± SEM, each dot represents a biologically independent observation.

## References

1. Abramson J, Anderson G. Thymic Epithelial Cells. Annu Rev Immunol. 2017;35:85–118.

2. Granadier D, Iovino L, Kinsella S, Dudakov JA. Dynamics of thymus function and T cell receptor repertoire breadth in health and disease. Seminars in Immunopathology. 2021.

3. Gruver AL, Sempowski GD. Cytokines, leptin, and stress-induced thymic atrophy. Journal of Leukocyte Biology. 2008;84(4):915–923.

4. Small TN, Papadopoulos EB, Boulad F, et al. Comparison of immune reconstitution after unrelated and related T-cell-depleted bone marrow transplantation: effect of patient age and donor leukocyte infusions. Blood. 1999;93(2):467–480.

5. Maury S, Mary JY, Rabian C, et al. Prolonged immune deficiency following allogeneic stem cell transplantation: risk factors and complications in adult patients. Br J Haematol. 2001;115(3):630–641.

6. Storek J, Joseph A, Espino G, et al. Immunity of patients surviving 20 to 30 years after allogeneic or syngeneic bone marrow transplantation. Blood. 2001;98(13):3505–3512.

7. Storek J, Gooley T, Witherspoon RP, Sullivan KM, Storb R. Infectious morbidity in long-term survivors of allogeneic marrow transplantation is associated with low CD4 T cell counts. Am J Hematol. 1997;54(2):131–138.

8. Maraninchi D, Gluckman E, Blaise D, et al. Impact of T-cell depletion on outcome of allogeneic bone-marrow transplantation for standard-risk leukaemias. Lancet. 1987;2(8552):175–178.

9. Kinsella S, Dudakov JA. When the Damage Is Done: Injury and Repair in Thymus Function. Frontiers in Immunology. 2020;11:1745.

10. Velardi E, Tsai JJ, van den Brink MRM. T cell regeneration after immunological injury. Nature Reviews Immunology. 2020.

11. Kimura T, Kambe T. The Functions of Metallothionein and ZIP and ZnT Transporters: An Overview and Perspective. Int J Mol Sci. 2016;17(3):336.

12. Cousins RJ, Liuzzi JP, Lichten LA. Mammalian zinc transport, trafficking, and signals. J Biol Chem. 2006;281(34):24085–24089.

13. Takagishi T, Hara T, Fukada T. Recent Advances in the Role of SLC39A/ZIP Zinc Transporters In Vivo. Int J Mol Sci. 2017;18(12).

14. Honscheid A, Rink L, Haase H. T-lymphocytes: a target for stimulatory and inhibitory effects of zinc ions. Endocr Metab Immune Disord Drug Targets. 2009;9(2):132–144.

15. Vallee BL, Falchuk KH. The biochemical basis of zinc physiology. Physiol Rev. 1993;73(1):79–118.

16. Hojyo S, Fukada T. Roles of Zinc Signaling in the Immune System. Journal of immunology research. 2016;2016:6762343–6762343.

17. Neldner KH, Hambidge KM. Zinc therapy of acrodermatitis enteropathica. N Engl J Med. 1975;292(17):879–882.

18. Ogawa Y, Kinoshita M, Shimada S, Kawamura T. Zinc in Keratinocytes and Langerhans Cells: Relevance to the Epidermal Homeostasis. J Immunol Res. 2018;2018:5404093.

19. Brummerstedt E. Animal model of human disease. Acrodermatitis enteropathica, zinc malabsorption. Am J Pathol. 1977;87(3):725–728.

20. Macdonald JB, Connolly SM, DiCaudo DJ. Think zinc deficiency: acquired acrodermatitis enteropathica due to poor diet and common medications. Arch Dermatol. 2012;148(8):961–963.

21. Hojyo S, Miyai T, Fujishiro H, et al. Zinc transporter SLC39A10/ZIP10 controls humoral immunity by modulating B-cell receptor signal strength. Proc Natl Acad Sci U S A. 2014;111(32):11786–11791.

22. Colomar-Carando N, Meseguer A, Company-Garrido I, et al. Zip6 Transporter Is an Essential Component of the Lymphocyte Activation Machinery. J Immunol. 2019;202(2):441–450.

23. Anzilotti C, Swan DJ, Boisson B, et al. An essential role for the Zn2+ transporter ZIP7 in B cell development. Nature Immunology. 2019.

24. Mitchell WA, Meng I, Nicholson SA, Aspinall R. Thymic output, ageing and zinc. Biogerontology. 2006;7(5-6):461–470.

25. Golden MN, Jackson A, Golden B. EFFECT OF ZINC ON THYMUS OF RECENTLY MALNOURISHED CHILDREN. The Lancet. 1977;310(8047):1057–1059.

26. Wong CP, Song Y, Elias VD, Magnusson KR, Ho E. Zinc supplementation increases zinc status and thymopoiesis in aged mice. J Nutr. 2009;139(7):1393–1397.

27. Wertheimer T, Velardi E, Tsai J, et al. Production of BMP4 by endothelial cells is crucial for endogenous thymic regeneration. Science Immunology. 2018;3(19).

28. Bogale A, Clarke SL, Fiddler J, Hambidge KM, Stoecker BJ. Zinc Supplementation in a Randomized Controlled Trial Decreased ZIP4 and ZIP8 mRNA Abundance in Peripheral Blood Mononuclear Cells of Adult Women. Nutr Metab Insights. 2015;8:7–14.

29. Coto JA, Hadden EM, Sauro M, Zorn N, Hadden JW. Interleukin 1 regulates secretion of zinc-thymulin by human thymic epithelial cells and its action on T-lymphocyte proliferation and nuclear protein kinase C. Proc Natl Acad Sci U S A. 1992;89(16):7752–7756.

30. Schmitt TM, de Pooter RF, Gronski MA, Cho SK, Ohashi PS, Zuniga-Pflucker JC. Induction of T cell development and establishment of T cell competence from embryonic stem cells differentiated in vitro. Nat Immunol. 2004;5(4):410–417.

31. Holmes R, Zuniga-Pflucker JC. The OP9-DL1 system: generation of T-lymphocytes from embryonic or hematopoietic stem cells in vitro. Cold Spring Harb Protoc. 2009;2009(2):pdb prot5156.

32. Dudakov JA, Mertelsmann AM, O‘Connor MH, et al. Loss of thymic innate lymphoid cells leads to impaired thymopoiesis in experimental graft-versus-host disease. Blood. 2017;130(7):933–942.

33. Hassan MN, Waller EK. GVHD clears the Aire in thymic selection. Blood. 2015;125(17):2593–2595.

34. Krenger W, Hollander GA. The immunopathology of thymic GVHD. Semin Immunopathol. 2008;30(4):439–456.

35. Barsanti M, Lim JM, Hun ML, et al. A novel Foxn1eGFP/+ mouse model identifies Bmp4-induced maintenance of Foxn1 expression and thymic epithelial progenitor populations. Eur J Immunol. 2017;47(2):291–304.

36. Schulkens IA, Castricum KC, Weijers EM, Koolwijk P, Griffioen AW, Thijssen VL. Expression, regulation and function of human metallothioneins in endothelial cells. J Vasc Res. 2014;51(3):231–238.

37. Hershfinkel M. The Zinc Sensing Receptor, ZnR/GPR39, in Health and Disease. Int J Mol Sci. 2018;19(2).

38. Fujie T, Segawa Y, Uehara A, et al. Zinc diethyldithiocarbamate as an inducer of metallothionein in cultured vascular endothelial cells. J Toxicol Sci. 2016;41(2):217–224.

39. Seandel M, Butler JM, Kobayashi H, et al. Generation of a functional and durable vascular niche by the adenoviral E4ORF1 gene. Proc Natl Acad Sci U S A. 2008;105(49):19288–19293.

40. Kinsella S, Evandy C, Cooper K, et al. Attenuation of homeostatic signaling from apoptotic thymocytes triggers a global regenerative response in the thymus. bioRxiv. 2020;[Preprint](August 31, 2020 [cited 08/23/2021)):DOI: 2020.2008.2031.275834.

41. Prasad AS. Impact of the discovery of human zinc deficiency on health. J Am Coll Nutr. 2009;28(3):257–265.

42. Kasana S, Din J, Maret W. Genetic causes and gene-nutrient interactions in mammalian zinc deficiencies: acrodermatitis enteropathica and transient neonatal zinc deficiency as examples. J Trace Elem Med Biol. 2015;29:47–62.

43. Xu Y, Wang M, Xie Y, et al. Activation of GPR39 with the agonist TC-G 1008 ameliorates ox-LDL-induced attachment of monocytes to endothelial cells. Eur J Pharmacol. 2019;858:172451.

44. Zhu D, Su Y, Zheng Y, Fu B, Tang L, Qin YX. Zinc regulates vascular endothelial cell activity through zinc-sensing receptor ZnR/GPR39. Am J Physiol Cell Physiol. 2018;314(4):C404–C414.

45. Sunuwar L, Medini M, Cohen L, Sekler I, Hershfinkel M. The zinc sensing receptor, ZnR/GPR39, triggers metabotropic calcium signalling in colonocytes and regulates occludin recovery in experimental colitis. Philos Trans R Soc Lond B Biol Sci. 2016;371(1700).

46. Clave E, Lisini D, Douay C, et al. A low thymic function is associated with leukemia relapse in children given T-cell-depleted HLA-haploidentical stem cell transplantation. Leukemia. 2012;26(8):1886–1888.

47. Monroe RJ, Seidl KJ, Gaertner F, et al. RAG2:GFP knockin mice reveal novel aspects of RAG2 expression in primary and peripheral lymphoid tissues. Immunity. 1999;11(2):201–212.

48. Alves NL, Huntington ND, Mention JJ, Richard-Le Goff O, Di Santo JP. Cutting Edge: a thymocyte-thymic epithelial cell cross-talk dynamically regulates intrathymic IL-7 expression in vivo. J Immunol. 2010;184(11):5949–5953.

49. Peukert S, Hughes R, Nunez J, et al. Discovery of 2-Pyridylpyrimidines as the First Orally Bioavailable GPR39 Agonists. ACS Med Chem Lett. 2014;5(10):1114–1118.

50. Reed NP, Henderson MA, Oltz EM, Aune TM. Reciprocal regulation of Rag expression in thymocytes by the zinc-finger proteins, Zfp608 and Zfp609. Genes Immun. 2013;14(1):7–12.

51. Han BY, Wu S, Foo CS, et al. Zinc finger protein Zfp335 is required for the formation of the naive T cell compartment. Elife. 2014;3.

52. Moore JB, Blanchard RK, Cousins RJ. Dietary zinc modulates gene expression in murine thymus: results from a comprehensive differential display screening. Proc Natl Acad Sci U S A. 2003;100(7):3883–3888.

53. King LE, Osati-Ashtiani F, Fraker PJ. Apoptosis plays a distinct role in the loss of precursor lymphocytes during zinc deficiency in mice. J Nutr. 2002;132(5):974–979.

54. Dudakov JA, Hanash AM, Jenq RR, et al. Interleukin-22 Drives Endogenous Thymic Regeneration in Mice. Science. 2012;336(6077):91–95.

55. Miyazono K, Maeda S, Imamura T. BMP receptor signaling: transcriptional targets, regulation of signals, and signaling cross-talk. Cytokine Growth Factor Rev. 2005;16(3):251–263.

56. Patel SR, Gordon J, Mahbub F, Blackburn CC, Manley NR. Bmp4 and Noggin expression during early thymus and parathyroid organogenesis. Gene Expr Patterns. 2006;6(8):794–799.

57. Gordon J, Patel SR, Mishina Y, Manley NR. Evidence for an early role for BMP4 signaling in thymus and parathyroid morphogenesis. Dev Biol. 2010;339(1):141–154.

58. Sharma D, Kanneganti TD. Inflammatory cell death in intestinal pathologies. Immunol Rev. 2017;280(1):57–73.

59. Lin PH, Sermersheim M, Li H, Lee PHU, Steinberg SM, Ma J. Zinc in Wound Healing Modulation. Nutrients. 2017;10(1).

60. Maret W. Zinc in Cellular Regulation: The Nature and Significance of “Zinc Signals”. Int J Mol Sci. 2017;18(11).

61. Nishida K, Hasegawa A, Yamasaki S, et al. Mast cells play role in wound healing through the ZnT2/GPR39/IL-6 axis. Sci Rep. 2019;9(1):10842.

62. Pongkorpsakol P, Buasakdi C, Chantivas T, Chatsudthipong V, Muanprasat C. An agonist of a zinc-sensing receptor GPR39 enhances tight junction assembly in intestinal epithelial cells via an AMPK-dependent mechanism. Eur J Pharmacol. 2019;842:306–313.

63. Shao Y, Wolf PG, Guo S, Guo Y, Gaskins HR, Zhang B. Zinc enhances intestinal epithelial barrier function through the PI3K/AKT/mTOR signaling pathway in Caco-2 cells. J Nutr Biochem. 2017;43:18–26.

64. Emri E, Miko E, Bai P, et al. Effects of non-toxic zinc exposure on human epidermal keratinocytes. Metallomics. 2015;7(3):499–507.

65. Chasapis CT, Loutsidou AC, Spiliopoulou CA, Stefanidou ME. Zinc and human health: an update. Arch Toxicol. 2012;86(4):521–534.

66. Bunting MD, Varelias A, Souza-Fonseca-Guimaraes F, et al. GVHD prevents NK-cell-dependent leukemia and virus-specific innate immunity. Blood. 2017;129(5):630–642.

67. Scarborough J, Mueller F, Arban R, et al. Preclinical validation of the micropipette-guided drug administration (MDA) method in the maternal immune activation model of neurodevelopmental disorders. Brain Behav Immun. 2020;88:461–470.

